# Sensitivity Analysis to Isolate the Effects of Proteases and Protease Inhibitors on Extracellular Matrix Turnover

**DOI:** 10.1101/2025.02.21.639501

**Authors:** Amirreza Yeganegi, Karla Robles, William J. Richardson

**Affiliations:** Department of Bioengineering, Clemson University, Clemson, SC; Ralpha E. Martin Department of Chemical Engineering, University of Arkansas, Fayetteville, AR

**Keywords:** protease, extracellular matrix, collagen turnover, network model, myocardial infarct

## Abstract

Matrix metalloproteinases (MMPs) are a family of proteases that drive degradation of extracellular matrix (ECM) across many tissues. MMP activity is antagonized by tissue inhibitors of metalloproteinases (TIMPs), resulting in a complex multivariate system with many MMP isoforms and TIMP isoforms interacting across a network of biochemical reactions – each with their own distinct kinetic rates. This system complexity makes it very difficult to identify which specific molecules are most responsible for driving ECM turnover in vivo and therefore the most promising therapeutic targets. To help elucidate the specific roles of various MMP and TIMP isoforms, we present a computational systems biology model of collagen turnover capturing all possible interactions between type I collagen, four different MMP isoforms (MMP-1, -2, -8, and -9), and three different TIMP isoforms (TIMP-1, -2, and -4). We used dye-quenched fluorescent collagen to monitor the degradation of collagen in the presence of various MMP+TIMP cocktails, and we then used these experimental data to fit hypothetical reaction system topologies in order to investigate their respective accuracies. We determined kinetic rate constants for this system and used post-myocardial infarct time courses of collagen, MMP, and TIMP levels to perform a parameter sensitivity analysis across the model reaction rates and predict which molecules and interactions are the important regulators of ECM in the infarcted heart. Notably, the model suggested that MMP degradation and inactivation terms were more important for driving collagen levels than TIMP interaction terms. In sum, this work highlights the need for systems-level analyses to distinguish the roles of various biomolecules operating with a complex system, prioritizes therapeutic targets for post-infarct cardiac remodeling, and presents a computational framework that can be applied to many other collagen-rich tissues.

## Introduction

Collagen is the most abundant structural protein in the human extracellular matrix (ECM), comprising one-third of the total protein in the human body^1^. Matrix metalloproteinases (MMPs) are zinc-dependent enzymes that participate in ECM degradation, and they cleave all structural components of the ECM. There are different subgroups of MMPs based on their substrate preferences, including, the collagenases (MMP-1, -8, and -13) that cleave fibrillar type collagens at a highly conserved site and the gelatinases (MMP-2 and -9) that degrade gelatins^2–4^. Tissue inhibitors of metalloproteinases (TIMPs) are specific inhibitors of MMPs which bind to active MMPs with 1:1 stoichiometry and control the activity of MMPs in tissues to help modulate the balance between collagen deposition and destruction^2,3,5^.

Collagen turnover is centrally involved in many diseases, including tissue fibrosis, cancer metastasis, wound healing, and myocardial infarction (MI)^6^. Collagen turnover depends on a highly regulated balance between collagens, MMPs, and TIMPs, but the diversity of matrix-MMP-TIMP interactions makes it very difficult to intuitively predict the effects of any individual matrix-, MMP-, or TIMP-targeting therapy. A computational model of collagen-MMP-TIMP could help us to better understand these interactions, enable us to systematically predict dynamic matrix turnover under diverse matrix, MMP, and TIMP levels, and eventually help us to use the model as a screening tool across patient-specific disease conditions.

Very few computational models have been previously published for matrix-MMP-TIMP interactions^7–9^, and all of them are focused on a small number of MMPs and TIMPs. For example, Karaggiannis and Popel’s computational model was limited to collagen I, MMP-2, MMP-14, and TIMP-2^8^. They included all known interactions between collagen I and two MMPs and one TIMP and represented each reaction as mass-action ordinary differential equations (ODEs) based on previously reported reaction rates. Vempati, Karaggiannis, and Popel also used ODEs to capture two MMPs and two TIMPs interactions^7^. These studies highlighted the ability of computational modeling to elucidate the possible interactions between substrate, MMPs and TIMPs and helped us understand the kinetics of these interactions. These findings also showed that there are possible unknown MMP-MMP and MMP-TIMP interactions which can make complex molecules that can be investigated using computational modeling^10^.

In this present study, we built a novel systems biology model using ODEs to capture mass-action kinetics of reaction between type I collagen, four different MMP isoforms, and three different TIMP isoforms. We used dye-quenched fluorescent collagen to monitor the degradation of collagen in the presence of various MMP+TIMP cocktails, fit theoretical reaction network topologies to these experimental data, and then conducted a parameter sensitivity analysis to rank the relative contributions of each MMP and TIMP reaction to collagen content in the heart after MI.

## Materials and Methods

### Experimental Data Collection

In order to experimentally test the effect of different MMP-TIMP combinations on collagen degradation, we prepared a 96-well plate with a range of different mixtures of Dye Quenched (DQ) collagen, MMPs, and TIMPs (Table 1). We used DQ collagen to monitor the proteolytic activity of different MMPs and MMP-TIMP mixtures. We used a large-scale system of nonlinear ODEs capturing the interactions of collagen I with four MMPs and three TIMPs to record reaction rates between these species. The DQ collagen labeled with fluorescein so heavily that the fluorescence signal is quenched in intact collagen. This substrate can be digested by most collagenases and gelatinases to yield highly fluorescent peptides. The increase in fluorescence signal is proportional to proteolytic activity and can be monitored using a fluorescence microplate reader.

**Table 1:** Experimental conditions spanning various MMP + TIMP combinations. All wells contained 5 µg of DQ collagen along with 0.05 µg of the indicated TIMPs and 0.1 µg of the indicated MMPs. All these combinations were then repeated with 0.2 µg of the indicated MMPs.

|  | MMP<br>1 | MMP<br>2 | MMP<br>8 | MMP<br>9 | MMPs<br>1+2+8 | MMPs<br>1+2+9 | MMPs<br>1+8+9 | MMPs<br>2+8+9 | MMPs<br>1+2+8+9 |
| --- | --- | --- | --- | --- | --- | --- | --- | --- | --- |
| TIMP 1 | + | + | + | + | + | + | + | + | + |
| TIMP 2 | + | + | + | + | + | + | + | + | + |
| TIMP 4 | + | + | + | + | + | + | + | + | + |
| TIMPS 1+2 |  |  |  |  | + | + | + | + | + |
| TIMPS 1+4 |  |  |  |  | + | + | + | + | + |
| TIMPS 2+4 |  |  |  |  | + | + | + | + | + |
| TIMPS 1+2+4 |  |  |  |  | + | + | + | + | + |

DQ collagen was purchased commercially (Thermo Fisher Scientific, Waltham, Massachusetts) and reconstituted to 1 mg/ml by adding 1 mL of deionized water (ddH2O) to the DQ substrate. The solution was then heated to 50°C and agitated in ultrasonic water bath for 5 minutes to facilitate dissolving. Recombinant human proMMP-1, proMMP-2, proMMP-8, and proMMP-9, solutions were purchased commercially (Enzo Life Sciences, Farmingdale, New York). ProMMPs were activated using provided activation protocol. ProMMP-1 was activated by 8µl of 0.5 mg/ml trypsin added to 100 µl of proMMP-1 and incubated for 30 minutes at 37°C. ProMMP-2 was activated by 2mM APMA (final concentration) for two hours at 37°C. ProMMP-8 was activated by 2µl of 0.5 mg/ml trypsin added to 100 µl of proMMP-1 and incubated for 30 minutes at 37°C. ProMMP-9 was activated by 20µl of 0.5 mg/ml TCPK-trypsin added to 100 µl of proMMP-9 and incubated for 30 minutes at 37°C. Recombinant protein TIMP-1, TIMP-2, and TIMP-4 were purchased in lyophilized form (ProSci, Poway, California) and reconstituted in ddH2O. The starting concentration of DQ collagen was 25 µg/ml for all conditions and the final volume of collagen-MMP-TIMP in each sample was 200 µl (5 µg of DQ collagen in every sample). The starting concentration of MMPs was either 0.1 µg or 0.2 µg, resulting in either 1:50 mass ratio of MMP:collagen or a 1:25 mass ratio. The starting concentration of TIMPs was held at 0.05 µg, resulting in either 1:2 mass ratio of TIMP:MMP or a 1:4 ratio. These collagen:MMP:TIMP mixtures were tested across a wide range of different MMP and TIMP isoforms in various combinations (Table 1).The degradation process was monitored using a fluorescent microplate reader in 37°C letting the proteases, inhibitors and substrate interact with each other for at least six hours. MMPs degrade DQ collagen and release fluorescein attached to the collagen. The product (degraded collagen) was measured using fluorescent microplate reader (BioTek, Synergy 4) with absorption maxima at 495 nm and fluorescence emission maxima at 515 nm.

### Computational Model Framework

A system of ordinary differential equations was used to mechanistically describe the MMP-substrate, MMP-TIMP, and MMP-MMP interactions. These models were constructed using mass action kinetics describing MMP binding and breakdown of substrates, along with TIMP binding and inhibition of MMPs. Our baseline model assumed one MMP binds and catalyzes collagen with associated *k*_*deg*_ (degradation) rates and one TIMP binds to one MMP with associated *k*_*inh*_ (inhibition) rates (Figure 1). Along with this baseline model, we tested the predictive accuracies of four additional model reaction topologies based on various complexities (Figure 1). The second model assumed one MMP binds to a substrate molecule with associated *k*_*on*_ and *k*_*off*_ rates and then cleaves the collagen to form degraded collagen and free enzyme with *k*_*deg*_ rate. TIMPs binding to MMPs also have *k*_*on*_ and *k*_*off*_ reaction rates instead of only degradation term. In the third model, MMP and TIMP inactivation terms were included (*k*_*inact*_). In the fourth model, MMP cannibalism terms were added which means that MMPs can interact with the same MMP in active and inactive form. In the final model, inactive MMPs and inactive TIMPs can be distracted by reacting with other TIMPs and MMPs as previously demonstrated in protease network interactions^10^.

**Figure 1:**
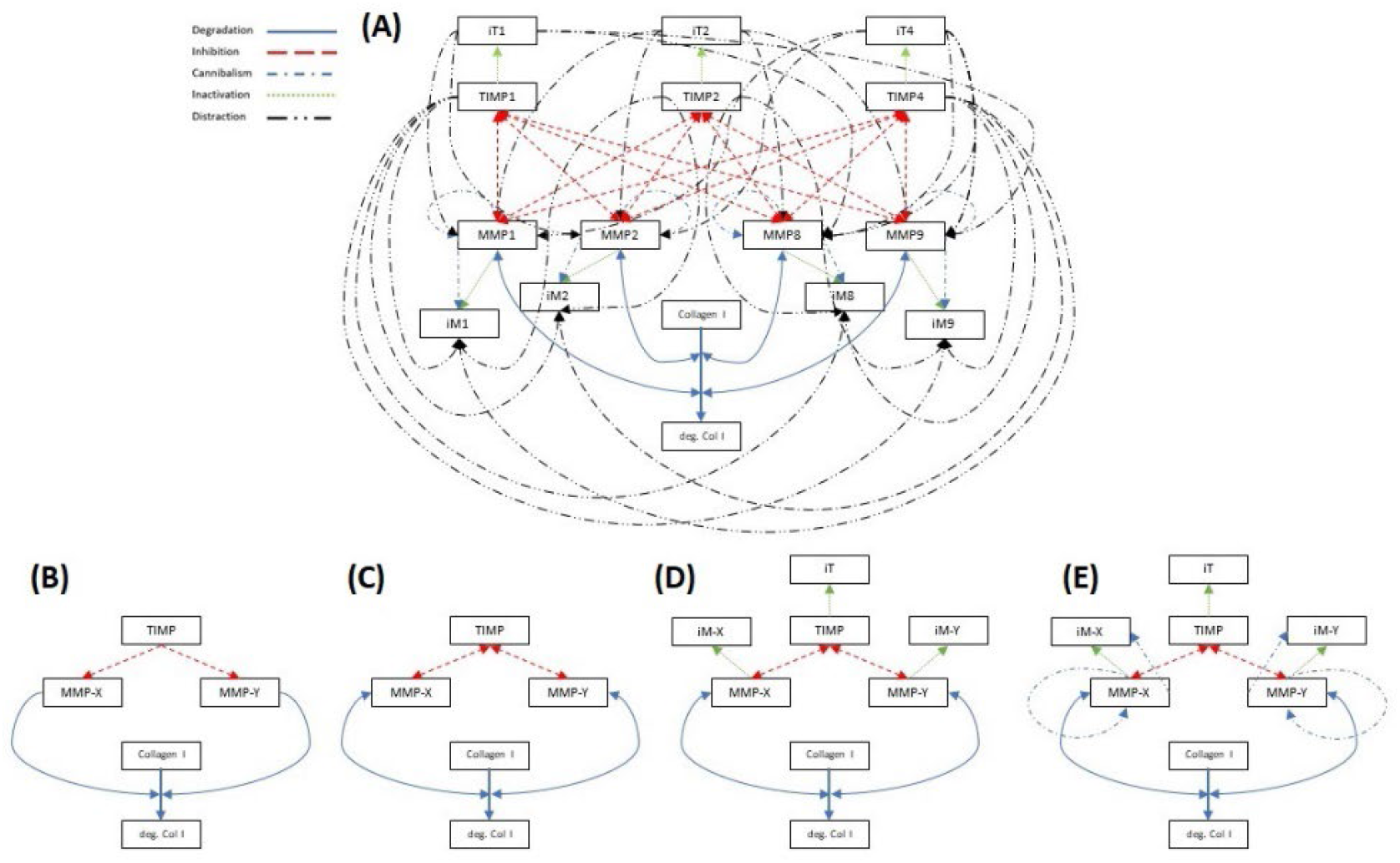
Schematic of different model reaction topologies. Blue solid lines show degradation of collagen by MMPs. Red dash lines show inhibition of MMPs by TIMPs. Blue, long dash dot lines show cannibalism of MMPs. Green dot lines show inactivation of MMPs and TIMPs. Black dash double dot lines show distraction of TIMP with inactive MMPs or distraction of MMPs by inactive TIMPs. A) The full model, containing MMP degradation, inhibition, and cannibalism, MMP and TIMP inactivation and MMP and TIMP distraction terms. B) Baseline model with one degradation and one inhibition term. C) Second model with “on” and “off” terms for MMP degradation and TIMP inhibition terms D) MMP and TIMP inactivation terms were added to the second model. E) MMP cannibalism terms were added. MMPs interact with the same MMP in active and inactive form.

The simplest model included 16 reaction rates (Figure 2B), and the most complicated model includes 163 reaction rates (Figure 2A). A simplified ODE system based on the most complicated model network is shown in equations 1-8 below. These equations capture the interactions of collagen I with one MMP and one TIMP. The complete interaction system follows these same forms for all possible MMP and TIMP isoforms.

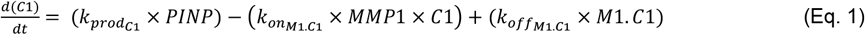

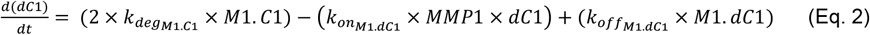

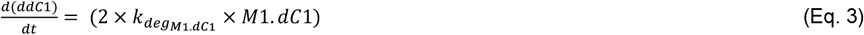

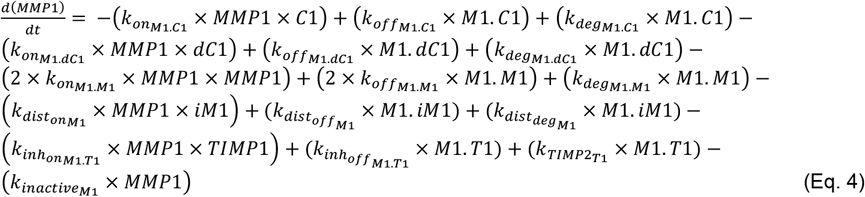

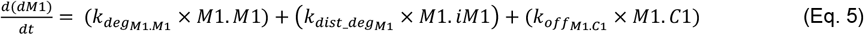

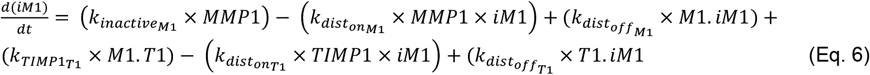

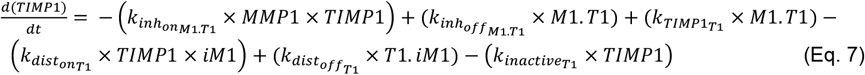

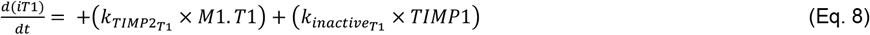

**Figure 2:**
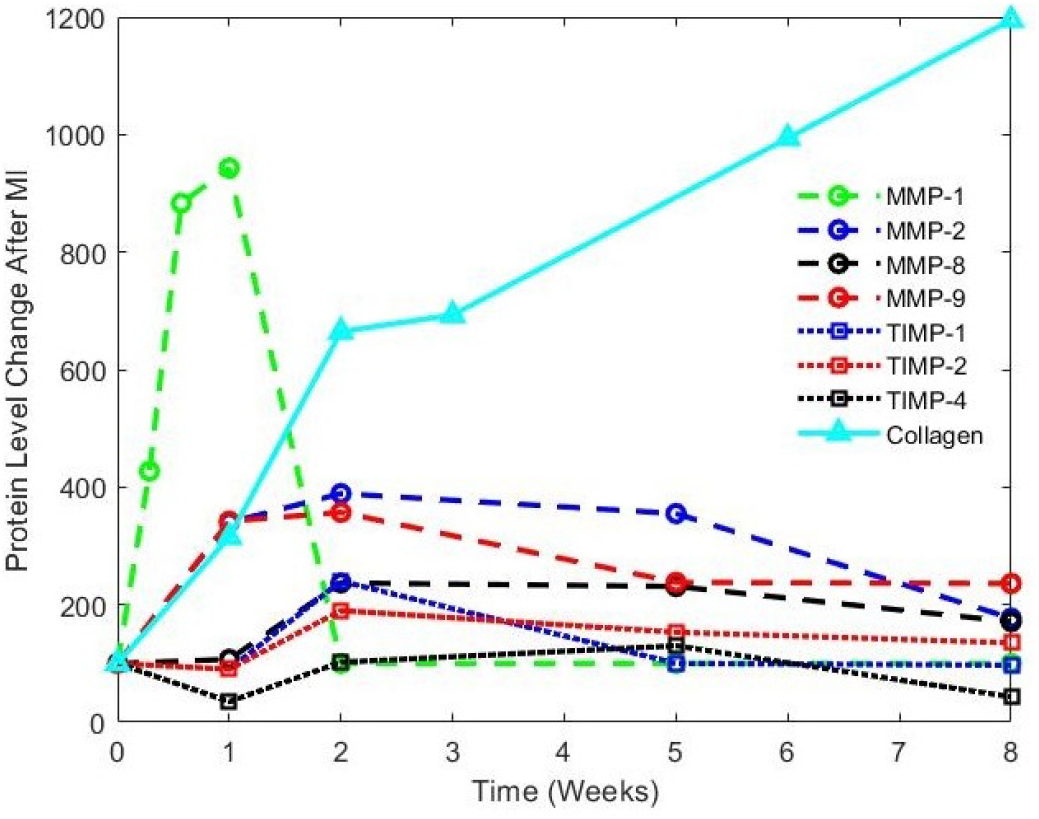
Levels of collagen, MMP, and TIMP in the heart measured following MI in mice shows that different MMPs are dominant in different time points.

The model kinetic rates were fit to experimental data by minimizing the difference between simulation predictions of degraded collagen levels and the experimentally measured levels in the conditions listed in Table 1. Experimental data were collected every 10 minutes for 6 hours, and thus, the simulations were also performed for 6 hours with time steps of 10 minutes generating same number of data point as the experimental data. The ODE system of collagen-MMP-TIMP interactions were approximated using the ode45 function in MATLAB (MathWorks), and a built-in genetic fitting algorithm was used to find an optimal (best-fit) combination of rate parameters. The objective function for the genetic algorithm was defined by adding sum of squared errors of experiment and simulation for all conditions described in Table 1. The genetic algorithm consisted of 50 generations, each with a population size of 100 sets of parameters. The initial constraints for the lower and upper bounds of kinetic parameters were set to 0 and 20 (*M* × *hr*^−1^ for bimolecular reaction coefficients;*hr*^−1^for unimolecular reaction coefficients) in an effort to avoid constraining the algorithm too tightly. After every generation, the top 20% of best-fitting parameter sets were used as “parents” for the next round of simulations; these sets were re-used in the next series of simulations; remaining parameter sets were generated by crossing the values contained in each set^13^. In order to reduce the likelihood of our model being trapped in local minimum, we repeated the same genetic algorithm 10 times using different initial values for parameters guesses, which created 10 different best-fit parameter combinations. All subsequent model simulations calculated ODE solutions using this ensemble of parameter combinations – in other words, final simulation results are the average of 10 different simulations, each of which relied on a different set of best-fit parameters.

### Parameter Sensitivity Analysis

Once our model was built and kinetic parameters were fit to our experimental data, we sought to test how strongly each parameter contributed to collagen turnover in an in vivo disease context, specifically, post-infarct scar tissue in the heart. We mined the literature to map post-MI time-courses of collagen, MMP, and TIMP levels in the heart in animal experiments^14,15^ (Figure 2). The concentration of collagen, MMPs and TIMPs varies at different time points after MI. We fed these concentrations at different time points as initial conditions into our model, and we then performed a parameter sensitivity analysis across the model reaction rates wherein we modulated each parameter in the network from 0.5x - 2x times their baseline values, one-at-a-time, and calculated the resulting changes in the remaining collagen concentration.

For each of the five models shown in Figure 1, we changed each reaction rate in the network from 0.5x - 2x times their baseline values and fit a linear function to 9 different collagen remaining values (0.5x 0.8x 0.9x 0.95x 1x 1.05x 1.1x 1.2x 2x). The mean of the slope of the 10 resulting lines for each kinetic rate parameter calculated from the ensemble of 10 different simulations was reported as the sensitivity of the model to that specific parameter. The same method was also performed for calculating the sensitivity of the model to the initial conditions^16^. The sensitivity analysis was repeated using inputs that match stimulation levels at 1-week, 2-weeks, 5-weeks, and 8-weeks post-MI in order to identify which species are the important regulators of ECM post-MI for early and late time points.

## Results

We incubated four different MMP isoforms and three different TIMP types with DQ collagen in different combinations to record time-courses of collagen degradation under various MMP-TIMP cocktails. Figure 3 shows the results of the final model comparison to experimental data for a subset of experimental conditions where collagen was treated with a single MMP in combination with each TIMP individually. Not surprisingly, different MMPs showed different degradation rates with MMP demonstrating the strongest proteolytic activity over MMP1, MMP8, and MMP9. Results also highlight that different TIMPs are more or less dominant for each MMP solution. As an example, TIMP-4 inhibits MMP-1 more effectively than the other TIMPs, while TIMP-1 inhibits MMP-2 more effectively.

**Figure 3:**
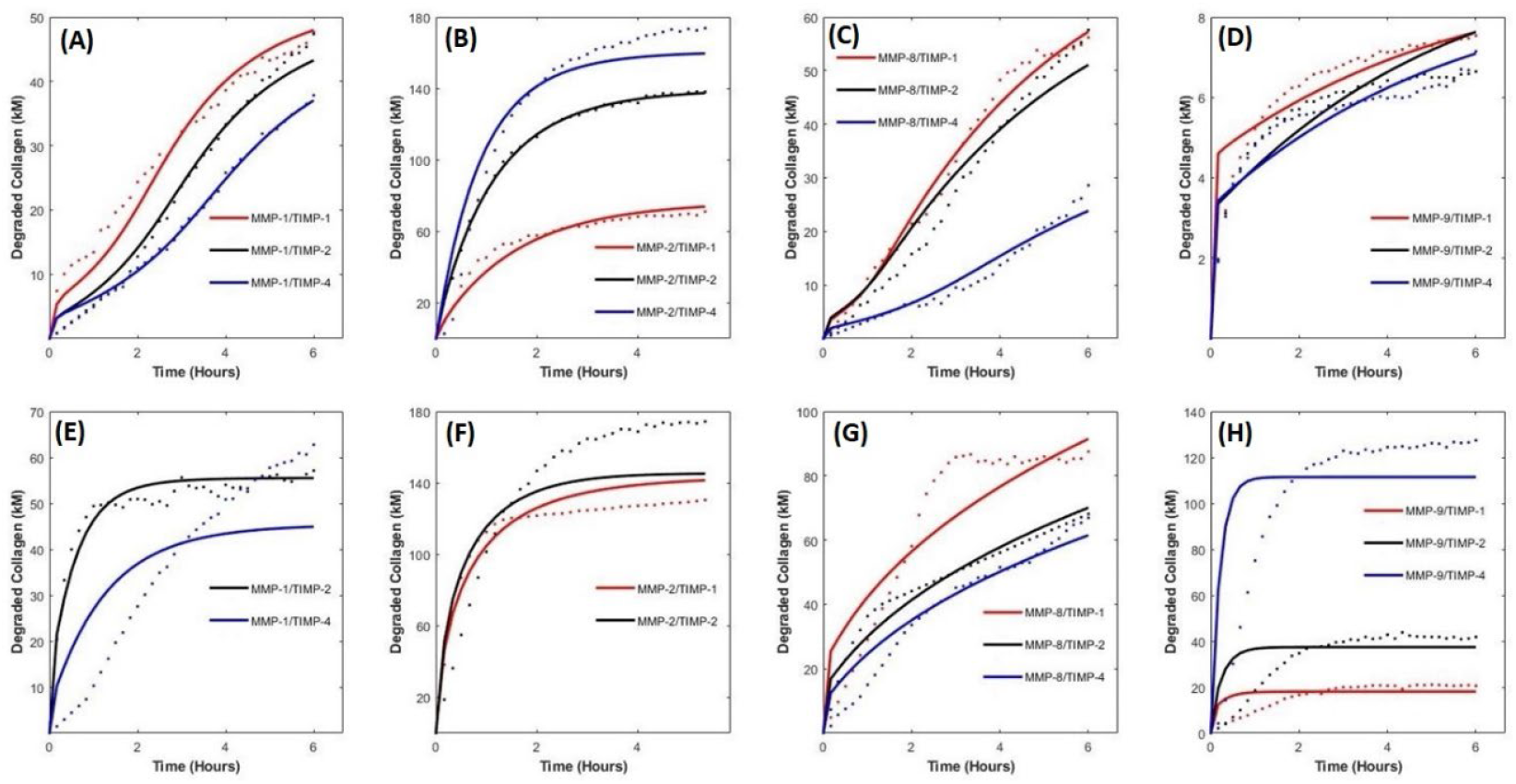
The final model results (solid lines) fitted to the experimental data (dots). Panels A-G show just a subset of conditions where collagen was treated with a single MMP in combination with different TIMPs. A) Low concentration MMP-1 and all TIMPs. B) Low concentration MMP-2 and all TIMPs. C) Low concentration MMP-8 and all TIMPs. D) Low concentration MMP-9 and all TIMPs. E) High concentration MMP-1 and all TIMPs. F) High concentration MMP-2 and all TIMPs. G) High concentration MMP-8 and all TIMPs. H) High concentration MMP-9 and all TIMPs.

In order to find the best model reaction topology to capture collagen turnover, we constructed five different models (Figure 1) and compared their abilities to match the experimental data via sum of squared errors (Figure 4). Using a genetic algorithm, each topology from simplest (topology 1) to most complex (topology 5) was fit to the experimental data 10 different times, resulting in 50 different best-fit parameter sets (ensemble of 10 sets for 5 different topologies). This comparison revealed substantial improvement from first model to the second model by replacing one step degradation with two step degradation (adding “on” and “off” terms to the collagen degradation kinetic). An additional improvement jump was seen between the second and third topology after adding inactivation terms to the model. Further adding complexity to the model from topology 3 to topologies 4 and 5 showed only modest reductions in model errors. Although the most complex model showed the best results (especially in low concentration cases) by capturing all possible interactions between species including cannibalism and distraction, the third model containing only degradation, inhibition, and inactivation terms was able to minimize the sum of squared errors to a comparable level to topology 4 and topology 5.

**Figure 4:**
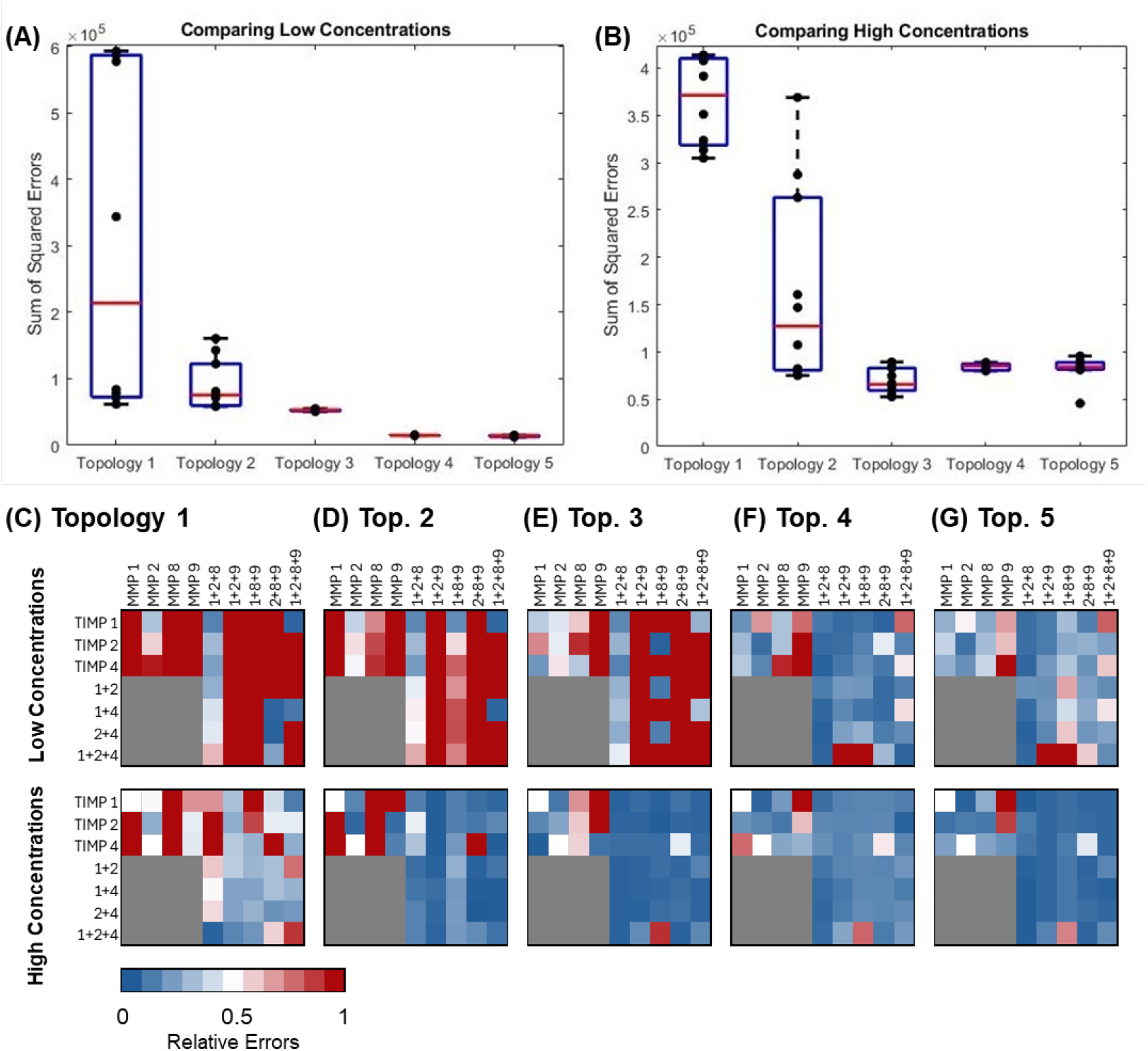
Simulation results (sum of squared error of all conditions) across five different topologies. Panels A-B show each dot as the error for a different best-fit parameter set compared to low MMP concentration data (A) or high MMP concentration data (B). Panels C-G show the relative error for each experimental condition fit by topology 1 (C) through topology 5 (G).

Once we finalized parameter fitting for the full model topology, we performed a parameter sensitivity analysis across the model variables at 1-week, 2-weeks, 5-weeks, and 8-weeks post-MI in order to identify which species are the most important regulators of ECM post-MI for both early and late time points. We modulated each reaction rate in the network from 0.5x - 2x times their baseline values, one at a time, ran the simulation for 6 hours, and calculated the resulting changes in collagen remaining concentration. The sensitivity to a parameter was then reported as the slope of the line fitted to collagen concentration changes relative to parameter value changes. This sensitivity analysis was repeated to identify the most important species of ECM post-MI that change the primary simulation output more drastically for both early and late time points. As expected, the MMP-collagen binding rates, MMP-collagen degradation rates, and MMP concentrations are all negatively correlated with collagen content across all time points, whereas the MMP-collagen unbinding rate, MMP inactivation rates, and initial collagen content values are all positively correlated with long-term collagen content across all time points (Figure 6). Unexpectedly, results show that protease inhibition by TIMPs seems to play only a minor role in collagen turnover post-MI as modulating concentration of TIMPs and the TIMP-MMP reaction rates had little effect on changes in collagen content. In contrast, modifying MMP concentrations or MMP-collagen reaction rates produced substantial changes in collagen content. These sensitivity values were largest for MMP2 and smallest for MMP8 and progressively increased from week 1 to week 8.

**Figure 6:**
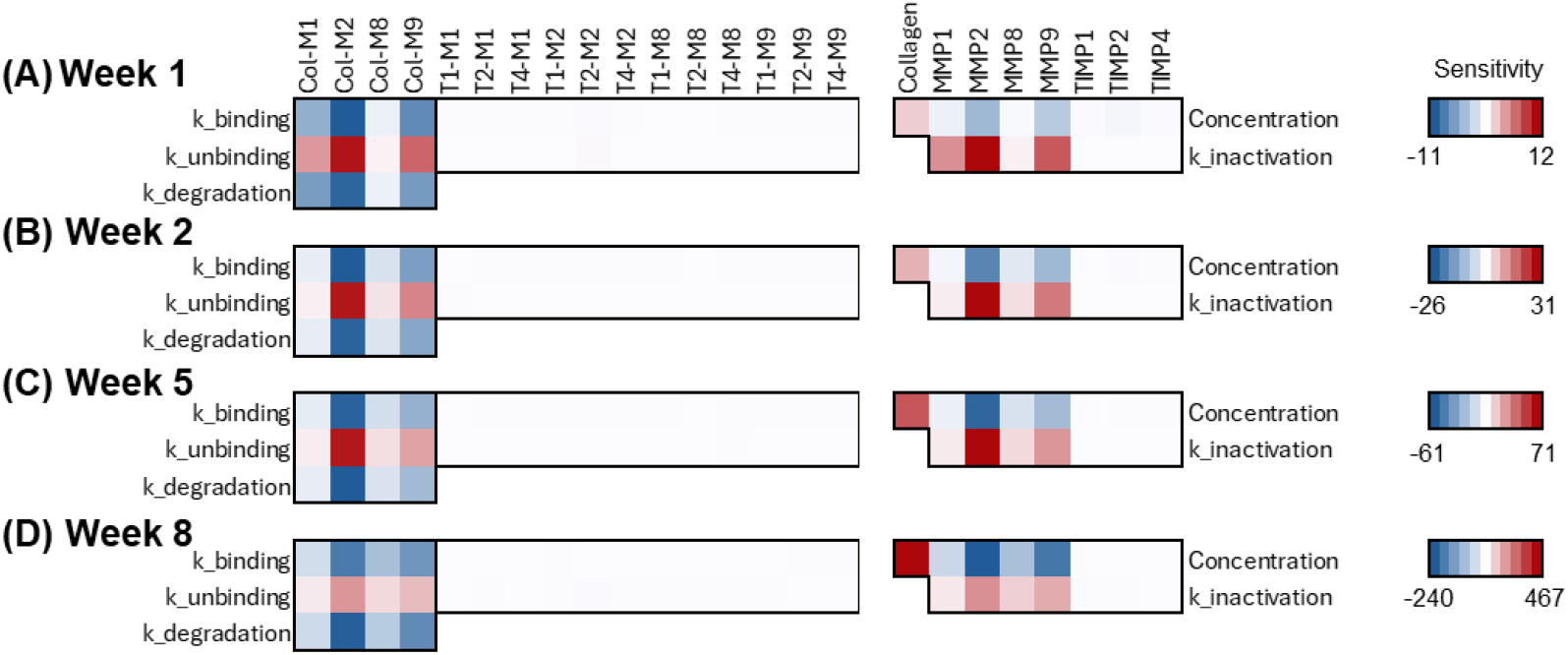
Sensitivity analysis for the rate parameters and initial conditions at 1-week, 2-weeks, 5-weeks and 8 weeks post-MI showing that the concentration of collagen is more sensitive to MMP-collagen interactions than it is to MMP-TIMP interactions. This analysis also shows that sensitivities progressively increase from week 1 to week 8, which coincides to progressive accumulation of collagen in the infarct scar.

## Discussion

The network of collagen-MMP-TIMP interactions is complicated by multifaceted crosstalk and nonlinear relationships that make it unclear which molecule or reaction is the most important in the cocktail mixtures within biological systems. We built a computational model that can integrate all multivariate interactions and capture the reaction parameters to investigate the kinetics of collagen-MMP-TIMP interactions. We fit the model to experimental data to better understand possible interactions and identify the key regulators of the system behavior, and we were able to successfully predict collagen degradation to for various MMP-TIMP cocktails in vitro. Simulating post-MI concentrations at different time points during heart attack healing suggested that MMPs (especially MMP2 and MMP9) play a larger role than TIMPs for determining collagen turnover. Future applications of this model could be helpful when used for predicting individual patients’ in-vivo contexts. Following MI, the concentration of ECM components, MMPs, and TIMPs change drastically, and the model can be helpful to predict collagen deposition to collagen degradation ratio as a means to monitor LV remodeling. Predicting collagen concentration after MI can help physicians develop potential treatments specifically modulated for each patient or for optimizing drug time-course strategies.

Of course, there are many limitations to this work. First, we must note the model parameters were trained using in vitro degradation data of DQ collagen molecules in a liquid suspension. In actual tissues in vivo, collagen molecules are assembled into a complex hierarchical organized structure that could modify MMP-collagen interaction parameters. In fact, the fluorescein tags that give DQ-collagen its fluorescent reporting capability could also affect collagen binding to MMPs in the mixture. Still, we believe this approach should be a reasonable approximation for the other MMP and TIMP interaction terms tested in our model topologies (e.g. MMP-TIMP binding, inactivation, cannibalism, and distraction rates). Another substantial limitation in our in vivo predictions is our neglection of mechanical tension effects on collagen-MMP interactions. Many past studies have established that mechanical stretching of collagen fibers can modulate their susceptibility to protease degradation and this strain-sensitivity can depend on the particular protease isoform^17-18^. Given that infarct scar tissue is subjected to dynamic mechanical stresses and strains in the heart, our sensitivity analysis could produce different rankings for various MMP and TIMP contributions if we were to incorporate mechanical-dependent reaction rates. This is an interesting question for future studies as there is growing motivation for mechano-dependent therapies in the infarcted heart^19-20^. Lastly, we must also acknowledge that scar tissue in the body is not made solely of type I collagen, but is itself a complex mileau of different extracellular matrix proteins including type III collagen, type IV collagen, elastin, fibronectin, periostin, and many others. Each of these matrix molecules is likely to have their own biochemical interaction rates with various MMPs, so a future comprehensive model of matrix turnover in the heart must incorporate many, many more reaction parameters. We hope the work presented in this study offers possible approaches for continuing to expand such a modeling framework. Ultimately, a large-scale model of matrix+protease+inhibitor interactions will enable an improved understanding of the molecular regulation of scar tissue as well as a therapeutic discovery tool to design patient-specific, temporal-specific, and location-specific interventions to control cardiac remodeling.

## Acknowledgments

We gratefully acknowledge funding support from the National Institutes of Health (HL144927 and GM121342).

